# Systematic assessment of GFP tag position on protein localization and growth fitness in yeast

**DOI:** 10.1101/360636

**Authors:** Dan Davidi, Uri Weill, Gat Krieger, Zohar Avihou, Ron Milo, Maya Schuldiner

## Abstract

While protein tags are ubiquitously utilized in molecular biology, they harbor the potential to interfere with functional traits of their fusion counterparts. Systematic evaluation of the effect of protein tags on localization and function would promote accurate use of tags in experimental setups. Here we examine the effect of Green Fluorescent Protein (GFP) tagging at either the N or C terminus of budding yeast proteins on localization and functionality. We use a competition-based approach to decipher the relative fitness of two strains tagged on the same protein but on opposite termini and from that infer the correct, physiological localization for each protein and the optimal position for tagging. Our study provides a first of a kind systematic assessment of the effect of tags on the functionality of proteins and provides step towards broad investigation of protein fusion libraries.

**Highlights:**

- Protein tags are widely used in molecular biology although they may interfere with protein function.
- The subcellular localization of hundreds of proteins in yeast is different when tagged at the N or the C terminus.
- A competition based assay enables systematic deciphering of correct tagging terminus for essential proteins.
- The presented approach can be used to derive physiologically relevant tagged libraries.

## Report

Protein tags are essential for a variety of assays in biology - from affinity tags for protein purification to fluorescence tags for visualization. However, tagging proteins comes at a price: fusion proteins are different from their native form and may suffer from impaired activity, reduced stability, loss of binding partners, wrong targeting, etc[1–4]. Often, the same tag may induce different phenotypes depending on where it appears on the protein. Most protein tags are added to one of the two termini of the polypeptide (carboxy terminus (C’) or amino terminus (N’)). However, with no a-priori knowledge, choosing the appropriate tagging terminus for a protein of interest requires trial and error.

Here we report a systematic approach suited for gauging the effect of a tag on global protein functionality. We use a Green Fluorescent Protein (GFP) tag as a test case and rely on a recent comparison made between two whole-genome libraries of strains, each encoding one protein fused to GFP at either the N’ [5] or C’ [6,7], In this comparison it was shown that 515 proteins in yeast are differentially localized when tagged in the opposing termini (Fig. 1A). While protein function can be impaired without displaying a mis-localization, it is clear that a difference in localization affects the capacity of a protein to function properly in a cellular context. Hence, we chose these proteins to test our method: which tagged terminus represents the physiologically relevant localization of these proteins?

**Figure 1:**
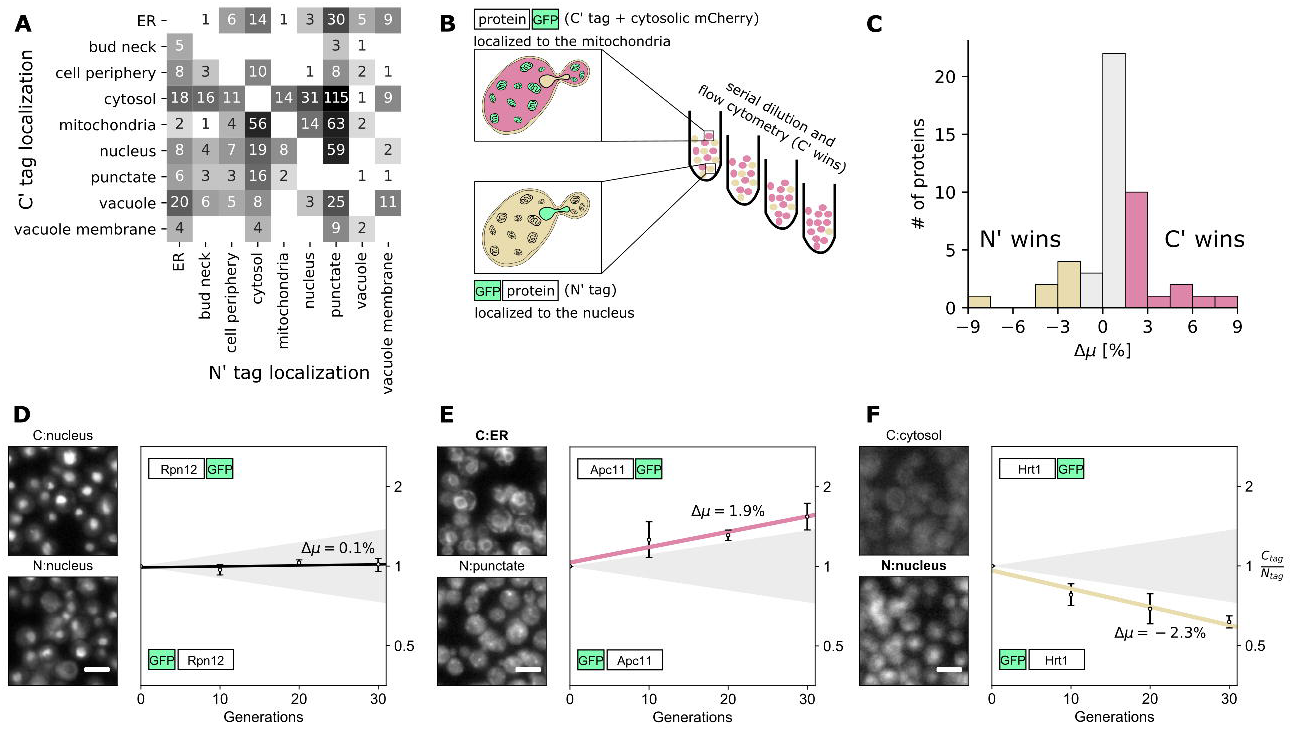
Uncovering the physiological subcellular localization of proteins in yeast. (**A**) Heatmap showing a comparison of localization assignments between the C’ tagged (y-axis;[6,7]) and N’ tagged GFP genome-scale library (x-axis;[5]). Altogether 515 proteins are differentially localized, representing about 10% of the entire collection of yeast proteins. Grayscale goes from white (least) to black (most) strains with altered localization (data from [5]) (**B**) Schematic representation of the pairwise competition approach. The C’ tagged library was genetically modified to express cytosolic mCherry, giving rise to the “red” phenotype, which in turn allows the quantitative measurement of population sizes of the two variants separately using flow cytometry on pooled mixed samples. Measurements were done every 24-hours for 4 days (about 30 generations; see Methods). (**C**) Distribution of the relative fitness (Δμ) for all essential proteins which are differentially localized (46 pairwise assays); yellow bars correspond to strains with Δμ<-1.5 and represent N’ winners, red bars are for Δμ>1.5 (C’ winners) and grey bars are within the noise range of the assay (determined empirically; see Methods). (**D, E & F**) Representative images showing proteins that (i) are localized to the same organelle and do not show a growth advantage with either tag (ii) localized to different places and show a growth advantage when harboring a C’ tag in comparison to an N’ tag and (iii) show a growth advantage with an N’ tag in comparison to a C’ tag. All measurements were done in triplicates; error bars represent the standard deviations; shaded triangles correspond to |Δμ|≤1.5; microscopy image scale-bars are 5μηη.

To systematically address whether an N’ or C’ tag better represents the correct cellular localization of a given protein, we established a pairwise competition approach that relies on the assumption that there would be a growth advantage to the strain carrying the correctly localized protein form (Fig. 1B). While it may theoretically be the case that mis-localization can give rise to a growth advantage, here we assume that this is not the norm. We hypothesized that such a difference could be easily monitored in essential proteins where even partial loss of the protein’s function inherently leads to a growth deficiency. Thus, to test this approach we focused on all the proteins in yeast that are both essential [8] and differentially localized (57 proteins, out of which 46 were successfully tested here; see Methods and Table S1 for further details). Flow cytometry was then used to infer the relative growth fitness difference (Δμ) for each pair of strains (N’ vs. C’ form) with identification of the fittest strain (as illustrated in Fig 1B; for full description of the assay see the Methods section).

A total of 21 proteins (out of the 46) showed a significant fitness difference between the two tagged forms (|Δμ|>1.5%; Fig. 1C); 14 cases where the C’ tagged form was superior and 7 cases where the N’ had an advantage. For example, Apc11, a catalytic core subunit of the Anaphase-Promoting Complex/Cyclosome (APC/C), showed an ER localization when C’ tagged as opposed to a punctate localization with the N’ tag (Fig. 1E), and had a growth advantage when C’ tagged, suggesting that this protein may serve as a connection between the ER and cell cycle regulation. Another example is Rsp5 that has an advantage when it is localized to the nucleus with the C’ tag while being in a punctate form when N’ tagged. This may suggest that either its SUMO ligase activity [9] is its essential function or that the control of multivesicular body (MVB) sorting [10] is also achieved through nuclear control (Table S1). An example of a N’ tag winner is Hrt1, a RING-H2 domain core subunit of multiple ubiquitin ligase complexes, that showed an advantage when localized to the nucleus with an N’ tag and mis-localized to the cytosol when C’ tagged (Fig 1F). Rpn12 is shown as a representative of a control, where both tagged forms are localized to the same organelle and the fitness of both strains is similar (Fig. 1D).

Notably, essential proteins that had a different localization but showed identical growth rate at our resolution level (the remaining 25 proteins; Table S1) may indicate that both localizations are tolerated (for example in dual localized proteins) or neither (both tags may cause mis-targeting of the protein). Comparison of each of the tagged variants to wild-type can help distinguish between the two cases, since, if both variants suffer from protein miss-localization, we expect the wild-type to be fitter than either. Here we used colony-size quantification to compare the fitness of N’ tagged variants to wild-type (Table S1). We found that out of the above 25 cases, where no significant fitness difference was found between the two forms, only 5 proteins had a significant reduction in colony-size relative to wild-type (mean=0.96, S.D=0.37 with a normal distribution according to the Shapiro-Wilk normality test), implying that in most cases where no superiority was observed, both localizations are tolerated. We are also aware that the presented approach may be more relevant for essential proteins, since, for non-essential proteins, the fitness difference between the mis- and well-localized variants may be too small to detect. However, many “non-essential” proteins become essential under specific conditions (different media and/or genetic backgrounds), and hence they could be included in tailored analyses. For example, peroxisomal biogenesis proteins become essential when cells are grown in fatty acids as a sole carbon source and mutants lacking mitochondrial genome become essential when yeast are forced to respire.

Our work suggests a systematic methodology to evaluate the effects of protein tags. The presented approach can readily be extended to study the effect of additional tags and therefore can be used to derive multiple physiologically relevant tagged libraries. In a similar manner, one can also test the effect of a given tag on the cellular *function* of a protein, by comparing the fitness of two variants that are localized to the *same* place. To conclude, we believe that our approach provides a useful tool to study the relationship between protein function and cellular fitness. Accounting for potential caveats of protein tags is essential for accurate understanding of cell biology. Such data are hence valuable for systematic, as well as for detailed, investigation of many questions in molecular biology.

## Methods

A total of 77 proteins were analyzed here (Table S1): 46 essential and differentially localized proteins (the study subset of study); 12 essential and similarly localized proteins; 9 non-essential and differentially localized proteins; and 10 non-essential and similarly localized proteins. For each protein, two strains were mixed in SD media such that one strain was tagged with GFP at the C’ of the protein of interest (taken from the genome wide C’ GFP yeast collection [7]) and the second strain was tagged with GFP at the N’ (taken from the N’ genome wide yeast collection *NATIVEpr-GFP [5]*). To allow optical separation between the strains, we included endogenous soluble mCherry in the C’ library strain (TEF2pr-mCherry tag was introduced into the URA3 locus; for more details see [7]). Cells were grown together for 24 hours, diluted 32-fold and then flow cytometry was used to monitor population sizes of the C’ and N’ tagged variants for 30 generations at 4 time points (every 24 hours for 4 days). To calculate the relative fitness difference (Δμ), we normalized the ratio between C’ and N’ by the ratio at day-zero to account for non-equal mixing of stains. Then, a linear regression model was fitted to the log of the ratio (y-axis in Fig. 1D,E&F) against the number of generations (x-axis in Fig. 1D,E&F). Δμ was calculated as the slope of the fit line and was positive if the C’ strain exhibited better fitness and negative if N’ was better. To evaluate the noise and sensitivity of the assay, 19 non-essential proteins were used; 9 with different localization and 10 with localization independent of the position of the terminus. A Δμ value of 1.5 (in absolute values) was considered to be a significant difference, just over the standard deviation in Δμ values of the 10 control strains which exhibited the same localization in both tag forms (Δμ=0.17 on average with S.D=1.42). For example, Δμ value of −1.8 means that N’ is 1.8% faster grower than the C’ strain. Flow cytometry was performed on the BD LSRII system (BD Biosciences). Fluorescent protein measurements were conducted with excitation at 488nm and emission at 525±25nm for GFP, excitation at 594nm and emission at 610±10nm for mCherry. The average number of cells analyzed was 20,000. Gating of +GFP-labeled population and +GFP+mCherry labeled population was done using a custom Matlab script; all measurements were done in triplicates. Downstream computational data processing was done using a custom Python script. We imaged the C’ and N’ GFP tagged strain arrays using a ScanR system (Olympus) as previously described [7], Images were acquired using a 60× air lens for GFP (excitation, 490/20 nm; emission, 535/50 nm), mCherry (excitation, 572/35 nm; emission, 632/60 nm), and brightfield channels. Images were transferred to ImageJ (1.51p Java1.8.0_144 (64-bit)), for slight, linear adjustments to contrast and brightness. Colony size quantification was done by plating yeast strains in 1536 format using a RoToR benchtop colony arrayer (PMID: 21877281) (Singer Instruments). Strains were grown overnight in 30°C and photographs of plates were analyzed for colony size using SGAtools [11]. Final Colony size score was calculated by dividing the colony size of a specific strain by the wild-type colony size from the same plate.

## Acknowledgments

We thank the Barkai lab for the kind help with generating the competition protocol. We thanks Naama Barkai and Einat Zalckvar for critical reading of the manuscript. This work was supported by the The Azrieli Institute of Systems Biology grant to DD and UW. Work in the Schuldiner lab is supported by an ERC CoG 646606 (Peroxisystem) and a VolksWagen foundation grant (93092). MS is an Incumbent of the Dr. Gilbert Omenn and Martha Darling Professorial Chair in Molecular Genetics.

